# Analysis of Ground Reaction Force Impulses During Uneven Walking for Young and Older Adults

**DOI:** 10.1101/2024.07.14.603453

**Authors:** Seyed Saleh Hosseini Yazdi

**Affiliations:** Department of Biomedical Engineering, Schulich School of engineering, University of Calgary, AB, Canada; Human Performance Laboratory, Faculty of Kinesiology, University of Calgary, AB, Canada

**Keywords:** Uneven walking, Ground Reaction Forces

## Abstract

Humans navigate various terrains by exerting forces to direct the Center of Mass (COM) and maintain balance. During walking, humans transition from one stance leg to the next by exerting impulses during the step-to-step transition. Studying these transition impulses may provide insight into how humans traverse uneven terrains. When walking speed increased (constant terrain amplitude), the average braking and propulsive impulses (posterior/anterior) increased comparably (−0.0270 m · s^−1^ · g^−1^ v^−1^versus 0.0252 m s^−1^· g^−1^ v^−1^). In the vertical direction, while the collision impulse remained constant, the push-off impulse declined by -0.0535 m · s^−1^ · g^−1^ · v^−1^. The interaction of age and speed also increased the collision impulse (0.0202 m · s^−1^ · g^−1^ · v^−1^). With the rise of terrain amplitude (constant speed), the braking and propulsive impulses rose by -0.0607 m · s^−1^ · g^−1^ · m^−1^ and 0.0701 m · s^−1^ · g^−1^ · m^−1^, respectively. Thus, we could infer that the propulsive impulse also contributed to the gait mechanical energy. While the collision impulse increased with terrain amplitude (0.1775 m · s^−1^ · g^−1^ · m^−1^) and the interaction of age and terrain amplitude (0.1058 m · s^−1^ · g^−1^ · m^−1^), the push-off impulse declined (−0.2700 m · s^−1^ · g^−1^ · m^−1^ and -0.1473 m · s^−1^ · g^−1^ · m^−1^). We also observed the push-off as a fraction of the total vertical impulse declined. Therefore, we detected a mechanical energy deficit in the step-to-step transition that must have been compensated for during the mid-flight phase. Considering the portion of push-off occurring after the subsequent heel-strike as delayed push-off, while it increased with walking speed, it declined with terrain amplitude. Thus, during uneven walking, the push-off exertion must have been interrupted, indicating the demand for further mechanical energy infusion after the transition.

## Introduction

Humans navigate various terrains that influence their walking characteristics. One of the walking objectives is to propel the body’s Center of Mass (COM) in the desired direction while preserving stability. Ground Reaction Forces (GRF) emerge from the interaction between the feet and the terrain, exerting impulses that alter human momentum. The resulting COM accelerations are pivotal for maintaining stability, balance, and efficient movement ^1^, while also facilitating energy conservation and conversion ^2^. Proprioceptive feedback from the body’s receptors concerning body position and motion are used for muscle contractions, aiding in balance maintenance and adjustments to walking patterns ^3^. Consequently, GRF analysis may serve as a valuable tool for gait assessment. As the step-to-step transition mechanical work is suggested to explain the majority of walking energetics ^4^, evaluating the associated impulses can enhance our understanding of momentum modulation, particularly in challenging walking conditions such as uneven terrain.

The vertical component of the GRF is predominantly studied due to its relatively large magnitude ^5,6^. It is hypothesized that the first peak of this component arises from weight acceptance and an increase in muscle forces, termed as the “collision” phase ^7,8^. The activation of the ankle plantar flexors, which push the body against the terrain, is suggested to contribute to the second peak ^1^. Upon initial contact, the Posterior/Anterior (P/A) force induces deceleration, reaching its maximum braking peak. As the stance progresses, this force shifts anteriorly, reaching its propulsive peak ^1,5^. While the Mediolateral (M/L) force influences body balance in the frontal plane ^1^, its analysis has been less frequent due to relatively high variabilities ^5^.

Studies investigating walking GRF have revealed the influence of walking speed on these forces. As walking speed rises, the magnitudes of the Posterior/Anterior (P/A) force peaks grow proportionally ^3,6^. Additionally, the magnitude of the first peak of the vertical force amplifies with velocity, although the growth of the second peak is comparatively smaller ^9^. It has been reported that in some cases, the amplitude of the second peak levels off. Conversely, lower walking speeds correspond to reduced Center of Mass (COM) accelerations, resulting in smaller vertical GRF peaks ^6^.

Experiments investigating the impact of aging on walking parameters have revealed distinct characteristics in older adults ^10^. The reported average preferred walking speed for older adults is approximately 1.0 m · s^−1^, while younger adults typical preferred walking speed ranges from 1.27 m · s^−1^ to 1.46 m · s^−1^ ^10,11^. Consequently, it’s reasonable to anticipate lower force peaks in the walking of older adults. Observations have indeed confirmed that vertical GRF peaks and troughs decrease with age. Yamada et al. ^12^ suggested that the reduced gait velocity in older adults is primarily attributed to decreased step length, leading to in a resultant GRF closer to vertical, thereby reducing Posterior/Anterior (P/A) force components. They further proposed that due to the reduced preferred speed, the vertical GRF peaks and troughs become smaller ^6^.

Vision also plays a crucial role in locomotion by contributing to body balance and orientation. Research indicates that visually impaired individuals also exhibit smaller GRFs and shorter step lengths at preferred walking speeds. While it is generally agreed that vision deprivation affects walking GRFs, results are somewhat varied. In many cases, subjects without vision exhibited a smaller vertical GRF second peak (push-off) and a less propulsive P/A peak ^13^, while in another study, a reduced first vertical GRF peak was observed ^14^.

When humans encounter terrain perturbations, their vertical GRF profile may alter. Aminiaghdam et al. ^15^ have reported that just before the perturbation, the second peak of vertical GRF decreases while the propulsive impulse increases. During and after the perturbation, they also reported that the second peak of vertical GRF increases, and the propulsive impulse decreases. They proposed that walking with a more crouched configuration observed in uneven walking leads to higher vertical GRF and loading rates during weight acceptance and a lower vertical GRF during the pre-swing phase. In another study, Aminiaghdam et al. ^16^ have suggested that by modeling the human leg as a spring and a damper, the damper skews the vertical GRF by increasing forces after touchdown and decreasing forces at toe-off, resulting in an early lift-off. Longer flight time during step-downs also suggested to result in a greater first peak of vertical GRF ^17^.

Human walking consists of discrete steps in which individuals transition from one stance leg to another. This transition is thought to involve mechanical work, starting with a pre-emptive push-off impulse by the trailing leg, followed by the collision of the leading leg’s heel strike ^7^. At any given velocity, if the nominal push-off is exerted, the subsequent collision has the same magnitude (Figure 1A) ^8^, while with a sub-optimal push-off, the magnitude of collision grows (Figure 1B) ^7^. Therefore, to sustain the gait, mid-flight positive mechanical energy inducement is necessary ^2,8^. This energy may come from the remainder of the push-off (delayed push-off ^18^) or from hip actuation ^19^. Since each step’s total positive mechanical work serves as a close proxy for the walking metabolic cost ^4^, changes in the magnitude of step-to-step transition impulses also influence the magnitude of the transition works ^20^.

**Figure 1:**
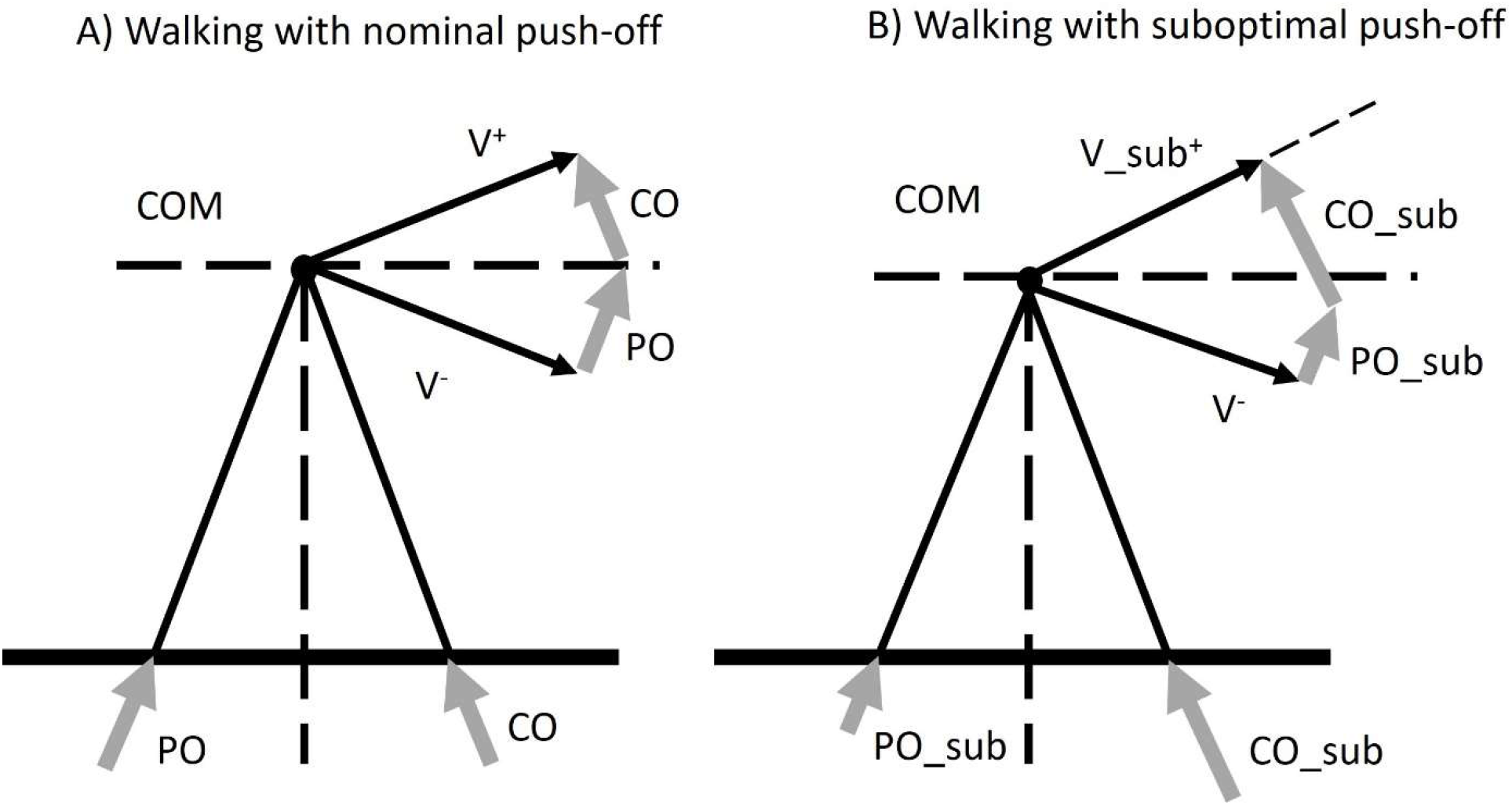
demonstration of powered simplest walking at the step-to-step transition: (A) in nominal walking the push-off is exerted pre-emptively that is followed by the collision. In nominal walking, the push-off and collision magnitudes are equal. (B) If suboptimal push-off is exerted, the magnitude of the following collision increases while the post transition velocity declines. Therefore, mid-flight energy compensation is needed.

Despite numerous studies exploring the effects of age, terrain, vision, and velocity on walking kinematics and kinetics ^21^, few have simultaneously considered these factors’ effects on GRFs and related them to energetics. Understanding the impact of these factors is crucial for studying walking pathology, stability, and balance. With these considerations, we hypothesize that uneven walking deviates GRF trajectories from the nominal profile, limiting push-off impulse exertion, which in turn yields larger collision impulses. Studying how individuals transition between steps reveals how walking terrain affects induced impulses and mechanical work performance, providing insight into how people adapt when faced with difficult conditions like walking on uneven terrain.

## Materials and Methods

To assess the effect of age on uneven terrain walking, we invited two groups of healthy adults to participate in our experiment: Young Adults (age: 27.69 ± 5.21, mass: 71.0 ± 6.16 kg, six females and four males) and Older Adults (age: 66.1 ± 1.44, mass: 77.34 ± 12.13 kg, six males and four females). The procedure was approved by the University of Calgary Conjoint Health Research Ethics Board, and subjects provided written informed consent before the experiments.

We modified a split belt instrumented treadmill (Bertec Corporation, Columbus, Ohio, USA) to accommodate uneven terrains by extending the supporting structure of the traveling section ^22^. Additionally, we fabricated uneven terrains by affixing construction foam blocks onto recreational treadmill belts. These foam blocks were arranged across the width of the belt, organized into sets measuring 0.3 m in length, with each set containing blocks of equal height. This configuration allowed for each foot to land flat on one set or at a slight angle across two adjacent sets. The heights of the sets were 0 m (Even), 0.019 m, 0.032 m, and 0.045 m, respectively. Each terrain was designated based on its maximum foam block height (Max_h, Figure 2A). The terrains were designed to be open-ended, to wrap them around the instrumented treadmill’s original belt when the tensioning rollers were released. When the terrains were closed using alligator lacing clips, the tensioning rollers were readjusted to their original position.

**Figure 2:**
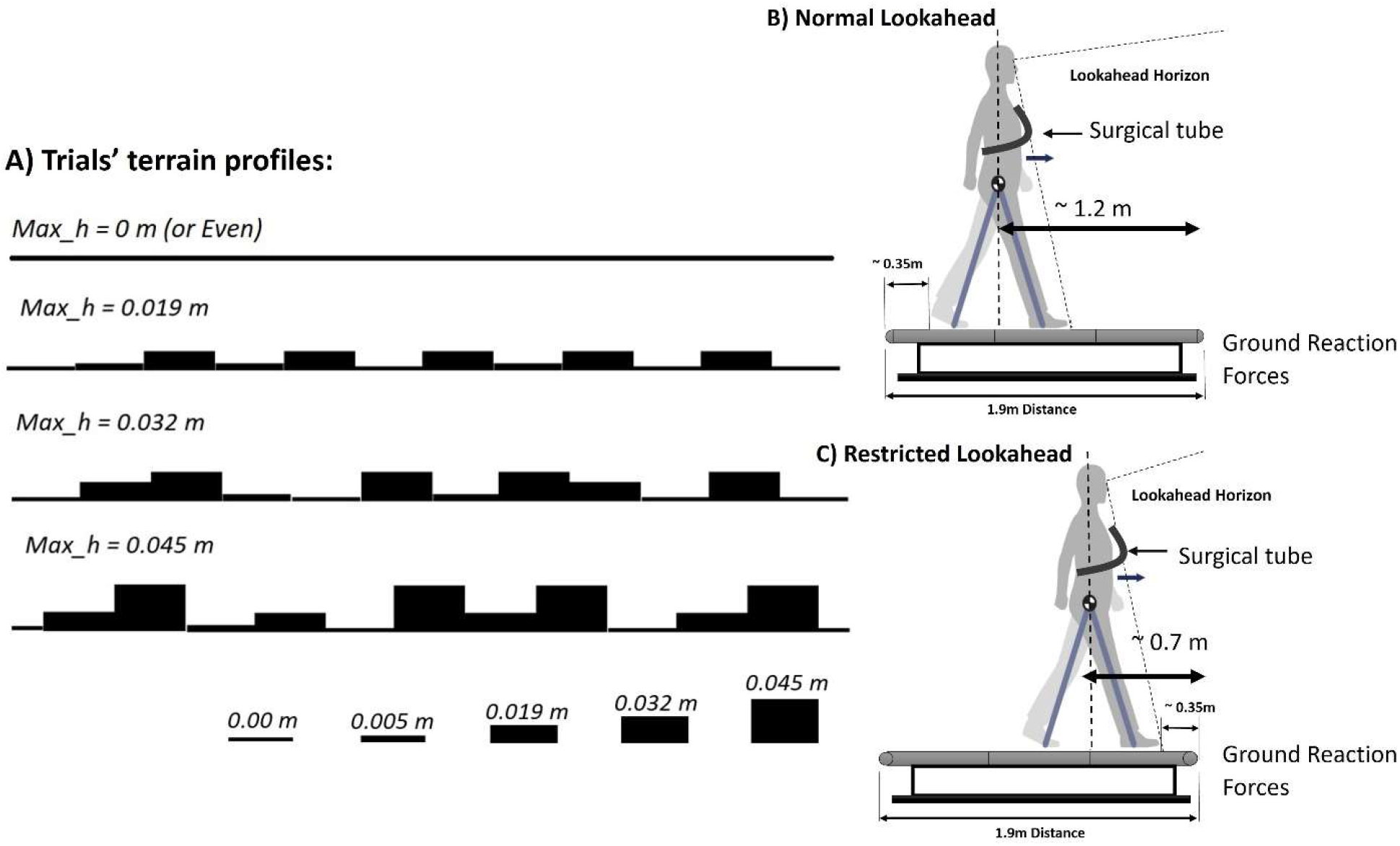
(A) Uneven walking experiments walking terrains profiles. The uneven terrains were wrapped around the original treadmill’s belts. (B) Normal lookahead at which subjects lookahead horizon was almost two steps, (C) Restricted lookahead in which the subjects’ lookahead horizon was limited.

To explore the influence of lookahead on uneven walking, we implemented two conditions: Normal Lookahead and Restricted Lookahead (Figure 2B & C). Under the Restricted Lookahead condition, participants were positioned approximately 0.7m away from the front end, instructing them to look forward ^22^. Conversely, in the Normal Lookahead condition, participants were situated 1.2 m behind the front end, allowing for approximately two step lengths of unobstructed view of the approaching terrain ^2^. Here, participants were encouraged to actively observe the approaching terrains while maintaining a normal walking. To ensure participants remained in the designated positions, we installed a flexible surgical tube across the treadmill. Participants were instructed to maintain gentle contact with the tube. Verbal feedback was provided if necessary. The sequence of trials was randomized (Table 1). For each trial, GRF data were collected for 60 seconds at a sampling rate of 960 Hz.

**Table 1:** The uneven experiment schedule: for Young (“YA” indicates normal lookahead and “ya” is for the restricted lookahead) and Older Adults (“OA”: is for normal lookahead and “oa” is for the restricted lookahead) uneven walking on different terrains and different speeds.

| Terrains Amplitudes (m) | 0 | 0.019 | 0.032 | 0.045 |
| --- | --- | --- | --- | --- |
| Trials Velocities (m/s) |  |  |  |  |
| 0.8 |  |  | oa-ya/ YA-OA |  |
| 1 |  |  | oa-ya/ YA-OA |  |
| 1.2 | oa-ya/ YA-OA | oa-ya/ YA-OA | oa-ya/ YA-OA | oa-ya/ YA-OA |
| 1.4 |  |  | oa-ya/ YA-OA |  |

The trial data was filtered using a third-order zero-phase Butterworth filter with a 10 Hz cut-off frequency, followed by time normalization. Subsequently, each participant’s GRF components in the vertical and P/A directions were normalized based on their individual weight^23^. Additionally, we computed vertical force collision and push-off impulses. The collision and push-off impulses were determined by integrating vertical forces from heel-strike to their first peaks and from their second peaks to toe-off, respectively ^24^. In the P/A direction, we calculated braking and propulsive impulses by integrating the negative and positive portions of the P/A force, respectively ^25^.

For the vertical force, we computed the delayed push-off by integrating the vertical force from the subsequent foot’s heel-strike to the end of the stance phase (double support). Additionally, we calculated the total vertical impulse by integrating the vertical force throughout the duration of the stance phase ^3,6^. We presented the total push-off (as fractions of the total vertical impulse) and delayed push-off (as a fraction of total push-off) variations with changes in walking velocity and terrain amplitude (Max_h, Figure 5A).

We conducted linear regression (Linear Mixed Effects Models, Statsmodels 0.15.0), with the impulses serving as the dependent variable. Trial walking velocities and terrain amplitudes (continuous variables), age group (categorical variable), and Normal Lookahead (versus restricted lookahead; categorical variable) were considered as independent variables. Regression coefficients and associated *p* values will be reported as measures to depict the relationship and significance (*p* < .05) between the variables of interest and the independent factors.

## Results

We investigated whether walking speed, terrain amplitude, age, and state of lookahead influence the trajectories of GRFs. Given that the step-to-step transition defines the COM work, serving as an indicator of walking energetics, we also analyzed their associated impulses variations under the specified conditions. We reported the rates of variation or offsets along with the measured ranges for young adults. Additionally, we reported the magnitude of variances if the measures for older adults differed significantly from those of young adults.

With visual inspection, we could recognize the changes in GRF components trajectories. Regardless of age and state of lookahead, the GRF components’ peaks appeared to change with terrain amplitude while their variations seemed to be more pronounced with the speed (Figure 3 and Figure 4). Walking at different speeds (Max_h = 0.032m, Table 2), the braking impulse magnitude increased significantly from -0.0235 m · s^−1^ · g^−1^to -0.0396 m · s^−1^ · g^−1^ (rate: - 0.0270 m · s^−1^ · g^−1^ · v^−1^, and 68.5% increase), while the propulsive impulse increased from 0.0282 m · s^−1^ · g^−1^ to 0.0421 m · s^−1^ · g^−1^ (rate: 0.0252 m · s^−1^ · g^−1^ · v^−1^, and 49.3% increase). The interaction of velocity and age further enhanced the rate of variation for the braking impulse by -0.0039 m · s^−1^ · g^−1^ · v^−1^ (14.4% rate increase). The braking and propulsion impulses did not change significantly with age. On the other hand, the propulsive impulse increased by 0.0013 m · s^−1^ · g^−1^ with the normal lookahead.

**Table 2:** The statistical result of linear regression analysis of the aggregate impulse data in P/A and vertical directions for different velocities over a constant terrain amplitude (Max_h = 0.032m). The young adults’ impulses with the normal lookahead are provided as a reference.

|  | P/A Braking Impulse |  |  |  |  | P/A Propulsive Impulse |  |  |  |
| --- | --- | --- | --- | --- | --- | --- | --- | --- | --- |
|  | Gains | P-Values | Velocities | YA<br>Normal<br>Lookahead |  | Gains | P-Values | Velocities | YA<br>Normal<br>Lookahead |
| Intercept* | -0.0028 | 4.99E-02 | 0.8 | -0.0235 | Intercept* | 0.0058 | 1.84E-08 | 0.8 | 0.0282 |
| Age | 0.0005 | 7.63E-01 | 1 | -0.0292 | Age | 0.0000 | 1.00E+00 | 1 | 0.0320 |
| Vision | -0.0002 | 6.29E-01 | 1.2 | -0.0341 | Vision* | 0.0013 | 1.74E-05 | 1.2 | 0.0370 |
| Velocity* | -0.0270 | 1.28E-230 | 1.4 | -0.0396 | Velocity* | 0.0252 | 1.55E-305 | 1.4 | 0.0421 |
| Velocity - Age* | -0.0039 | 1.84E-02 |  |  | Velocity - Age | 0.0000 | 1.00E+00 |  |  |
|  | Collision Impulse |  |  |  |  | Push-Off Impulse |  |  |  |
|  | Gains | P-Values | Velocities | YA<br>Normal<br>Lookahead |  | Gains | P-Values | Velocities | YA<br>Normal<br>Lookahead |
| Intercept* | 0.1062 | 6.74E-154 | 0.8 | 0.1088 | Intercept* | 0.1580 | 2.08E-263 | 0.8 | 0.1165 |
| Age | 0.0068 | 9.69E-02 | 1 | 0.1036 | Age* | -0.0131 | 2.50E-04 | 1 | 0.0997 |
| Vision* | -0.0036 | 9.29E-04 | 1.2 | 0.1018 | Vision | -0.0013 | 3.94E-01 | 1.2 | 0.0921 |
| Velocity | 0.0025 | 3.16E-01 | 1.4 | 0.1018 | Velocity* | -0.0535 | 2.09E-56 | 1.4 | 0.0888 |
| Velocity - Age* | 0.0202 | 1.29E-05 |  |  | Velocity - Age | -0.0036 | 5.94E-01 |  |  |

**Figure 3:**
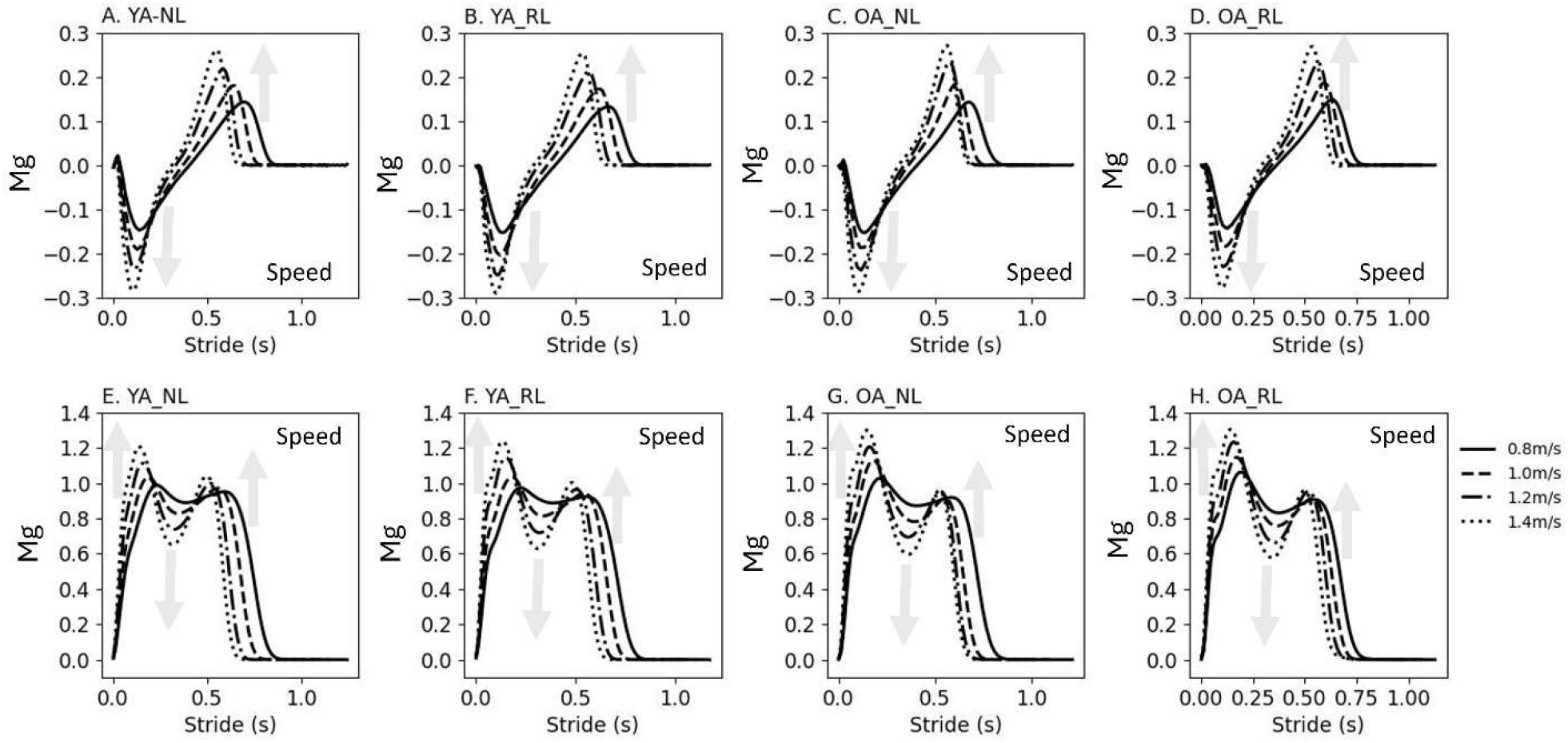
The average of ground reaction force (GRF) trajectories in P/A (top row: A. to D.) and vertical directions (bottom row: E. to H.) during uneven walking at different velocities (Max_h = 0.032m) for young (YA) and older adults (OA) with normal (NL) and restricted lookaheads (RL).

**Figure 4:**
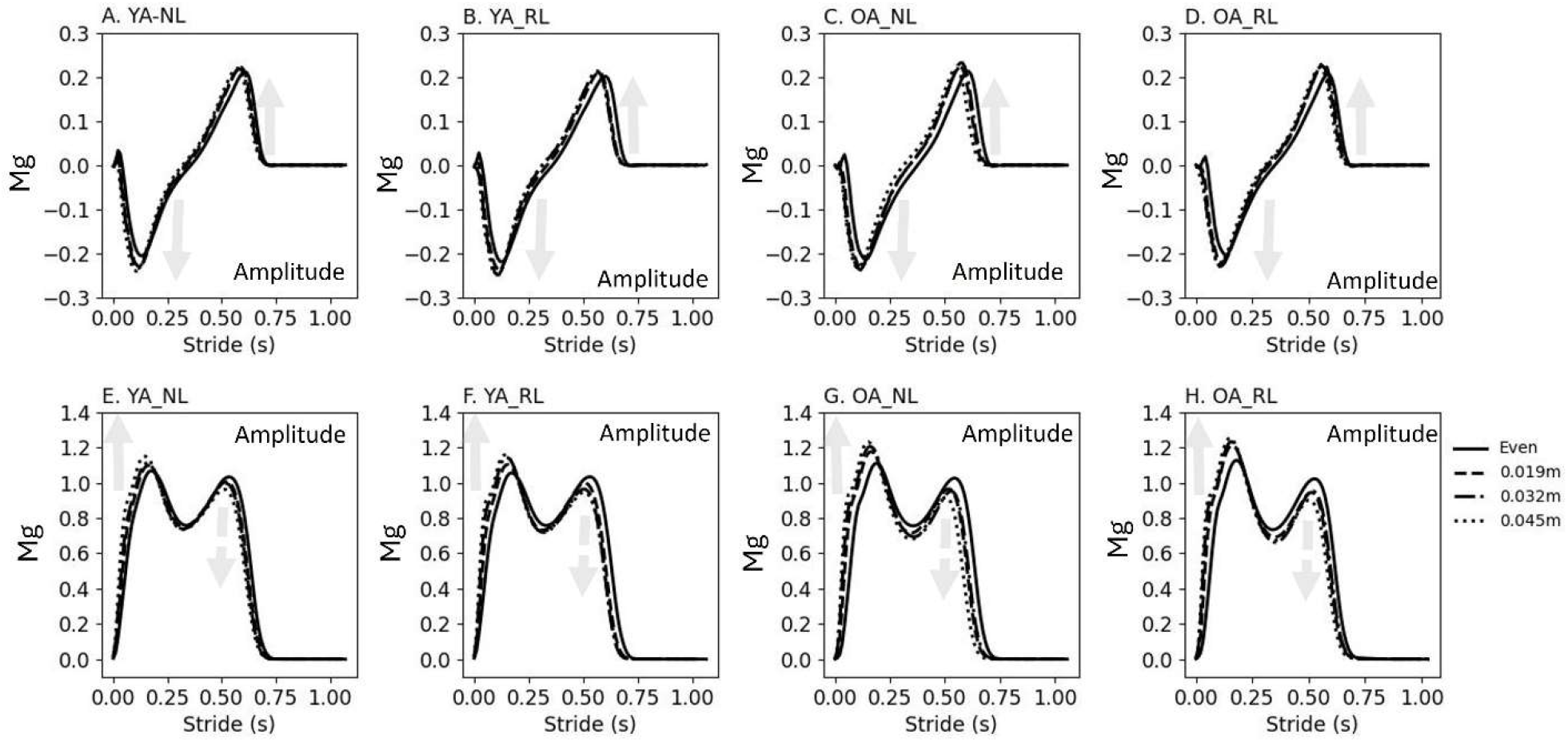
The average of ground reaction force (GRF) trajectories in P/A (top row; A. to D.) and vertical directions (bottom row: E. to H.) during uneven walking on different terrain amplitudes at a constant velocity (V=1.2 *m* · *s*^−1^) for young (YA) and older adults (OA) with normal (NL) and restricted lookaheads (RL).

The collision impulse only increased significantly with the interaction of velocity and age. For older adults, the collision impulses increased from 0.1082 m · s^−1^ · g^−1^ to 0.1162 m · s^−1^ · g^−1^ (rate: 0.0202 m · s^−1^ · g^−1^ · v^−1^, and 7.4% increase). With the normal lookahead, the collision impulse reduced by -0.0036 m · s^−1^ · g^−1^ (offset). The push-off impulse also declined significantly by age (offset: -0.0131 m · s^−1^ · g^−1^) and walking velocity (rate: -0.0535 m · s^−1^ · g^−1^ · v^−1^). For young adults, push-off impulse ranged from 0.1165 m · s^−1^ · g^−1^to 0.0888 m · s^−1^ · g^−1^ (23.8% decline).

Walking with a constant speed (v = 1.2 m · s^−1^) over different terrain amplitudes (Table 3), the braking impulse magnitude only increased by the training amplitude from -0.0309 m · s^−1^ · g^−1^ to -0.0342 m · s^−1^ · g^−1^ (rate: -0.0607 m · s^−1^ · g^−1^ · m^−1^, 10.7% rise), whereas the propulsive impulse increased with the normal lookahead (offset; 0.0013 m · s · g), and terrain amplitude from 0.0348 m · s^−1^ · g^−1^ to 0.0376 m · s^−1^ · g^−1^ (rate: 0.0701 m · s^−1^· g^−1^ · m^−1^, 8.0% increase).

**Table 3:**
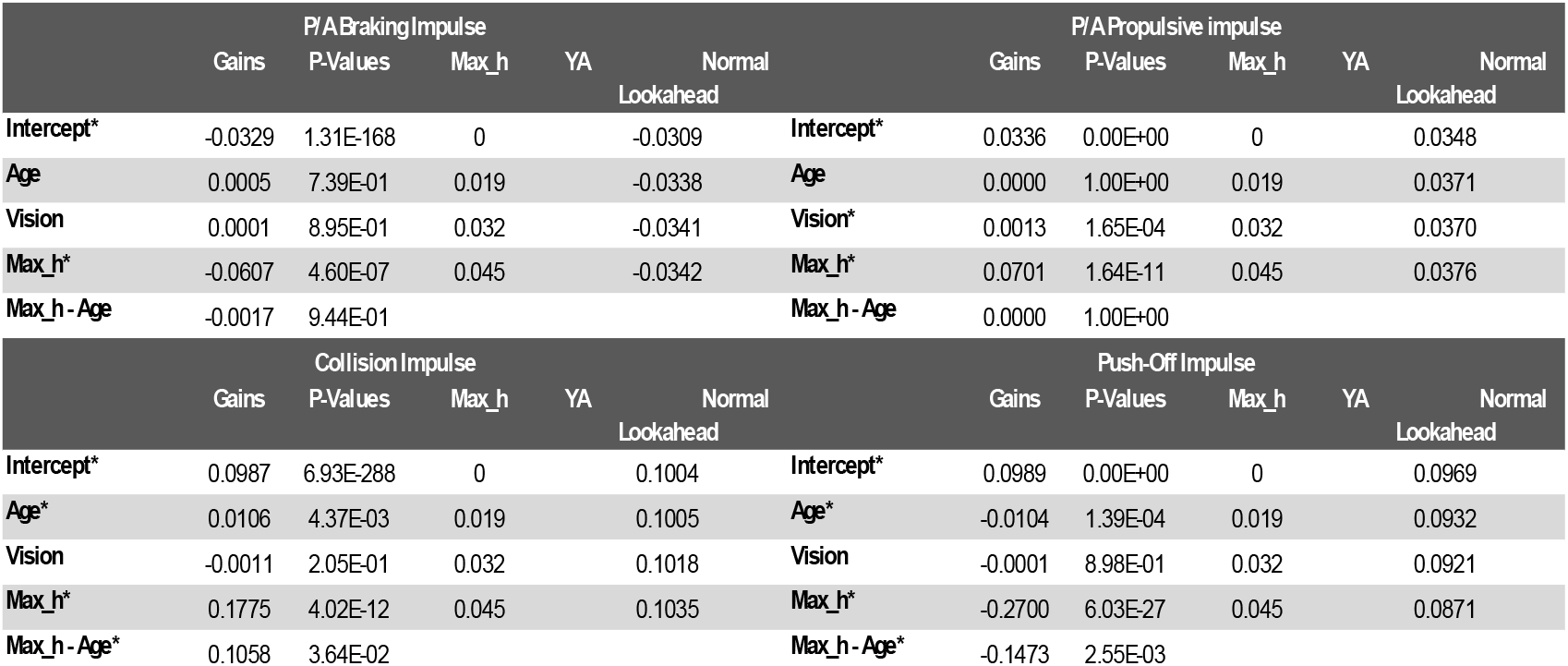
The statistical result of linear regression analysis of the aggregate impulse data in P/A and vertical directions for different terrain amplitudes with constant walking speed (v = 1.2 *m* · *s*^−1^). The young adults’ impulses with the normal lookahead are provided as a reference.

|  | P/A Braking Impulse |  |  |  |  | P/A Propulsive impulse |  |  |  |
| --- | --- | --- | --- | --- | --- | --- | --- | --- | --- |
|  | Gains | P-Values | Max_h | YA<br>Lookahead |  | Gains | P-Values | Max_h | YA<br>Lookahead |
| Intercept* | -0.0329 | 1.31E-168 | 0 | -0.0309 | Intercept* | 0.0336 | 0.00E+00 | 0 | 0.0348 |
| Age | 0.0005 | 7.39E-01 | 0.019 | -0.0338 | Age | 0.0000 | 1.00E+00 | 0.019 | 0.0371 |
| Vision | 0.0001 | 8.95E-01 | 0.032 | -0.0341 | Vision* | 0.0013 | 1.65E-04 | 0.032 | 0.0370 |
| Max_h* | -0.0607 | 4.60E-07 | 0.045 | -0.0342 | Max_h* | 0.0701 | 1.64E-11 | 0.045 | 0.0376 |
| Max_h - Age | -0.0017 | 9.44E-01 |  |  | Max_h - Age | 0.0000 | 1.00E+00 |  |  |
|  | Collision Impulse |  |  |  |  | Push-Off Impulse |  |  |  |
|  | Gains | P-Values | Max_h | YA<br>Lookahead |  | Gains | P-Values | Max_h | YA<br>Lookahead |
| Intercept* | 0.0987 | 6.93E-288 | 0 | 0.1004 | Intercept* | 0.0989 | 0.00E+00 | 0 | 0.0969 |
| Age* | 0.0106 | 4.37E-03 | 0.019 | 0.1005 | Age* | -0.0104 | 1.39E-04 | 0.019 | 0.0932 |
| Vision | -0.0011 | 2.05E-01 | 0.032 | 0.1018 | Vision | -0.0001 | 8.98E-01 | 0.032 | 0.0921 |
| Max_h* | 0.1775 | 4.02E-12 | 0.045 | 0.1035 | Max_h* | -0.2700 | 6.03E-27 | 0.045 | 0.0871 |
| Max_h - Age* | 0.1058 | 3.64E-02 |  |  | Max_h - Age* | -0.1473 | 2.55E-03 |  |  |

The collision impulse increased significantly with age (offset; 0.0106 m · s^−1^ · g^−1^). The collision impulses ranged from 0.1004 m · s^−1^ · g^−1^ to 0.1035 m · s^−1^ · g^−1^ (rate: 0.1775 m · s^−1^ · g^−1^ · m^−1^, and 3.1% rise). The interaction of terrain amplitude and age further enhanced the collision increase rate by 0.1058 m · s^−1^ · g^−1^ · m^−1^ (59.6% increase). On the other hand, push-off declined with age (offset: -0.0104 m · s^−1^ · g^−1^) and terrain amplitude (rate; -0.2700 m · s^−1^ · g^−1^ · m^−1^). The push-off reduced from 0.0969 m · s^−1^ · g^−1^ to 0.0871 m · s^−1^ · g^−1^ (11.3 % decline) as terrain amplitude increased. Additionally, the push-off decreased at a faster rate with the interaction of terrain amplitude and age. The additional decline rate was -0.1473 m · s^−1^ · g^−1^ · m^−1^ (54.6% faster decrease).

With the normal lookahead, the delayed push-off increased with the walking speed from 0.80 to 0.87 of the total push-off impulse (8.75% increase). With the restricted lookahead, the delayed push-off was offset upward by 0.04 of the total push-off. The delayed push-off only declined with the train amplitude from 1.0 to 0.82 of the total push-off (18.0% decline, Figure 5 – B to E).

**Figure 5:**
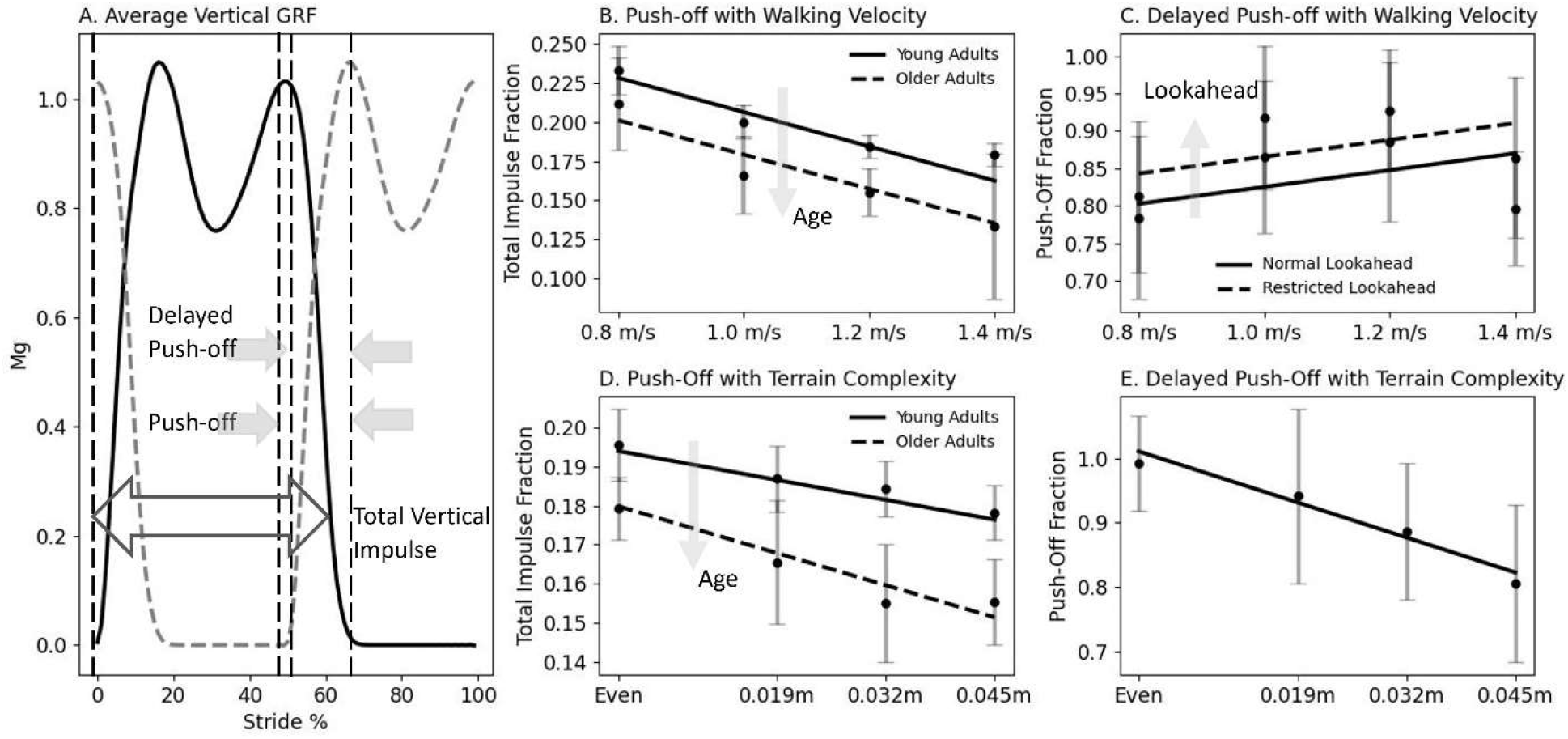
(A) Total vertical GRF impulse is the time integration of the vertical component of GRF over the stance phase. The Push-off impulse is assumed to start from the vertical GRF second peak and finishes by the stance phase end. The delayed push-off is the portion of the push-off impulse that starts when the leading leg heel-strike occurs while the trailing leg is still in contact with the ground. (B) the variation of push-off for young and older adults as fractions of total vertical impulse with walking velocity change. (C) the variation of delayed push-off as a fraction of total push-off with normal and restricted lookaheads. (D) the variation of delayed push-off for young and older adults with terrain amplitude increase. (E) the variation of delayed push-off as a fraction of total push-off with terrain amplitude rises.

## Discussion

Ground reaction forces result from the interaction between human legs and the walking terrain. As walking involves the alternation of stance legs, where mechanical energy is dissipated ^4^, the exerted impulses and associated mechanical works energize the step-to-step transition. Thus, it’s reasonable to infer that the way in which feet are planted and their timing might influence the trajectory of the GRFs and transition impulses. In this study, we aimed to investigate how GRF component impulses vary with walking velocity, terrain amplitude, age, and state of lookahead. We anticipated detecting variations in GRF impulses with the selected trial parameters.

Furthermore, we hypothesized that the imposed challenges in our trials would limit the magnitude of push-off, coinciding with an increase in collision impulse magnitude.

The P/A braking impulses partially dissipate the COM mechanical power during the collision phase ^25^. The vertical GRF collision impulse independence from velocity may suggest that the vertical GRF at early stance is solely determined by the walker’s weight, while the P/A braking impulse is proportional to the walker’s velocity. The decrease in vertical GRF collision impulse with normal lookahead (vision) also suggests that humans tend to choose appropriate foot placements to maximize the conversion of potential to kinetic energy ^2^ and thereby require less active work for the step-to-step transition. The increase in collision with the interaction of age and velocity in both P/A and vertical directions may indicate that with higher velocity, older adults may experience difficulty in regulating their foot placement properly, resulting in increased step-to-step dissipations.

The propulsive component of the P/A impulse, which contributes to the walking push-off, can also be assumed to be proportional to walking velocity. The similar rate of variation in propulsive impulse compared to braking impulse (0.0252 m · s^−1^ · g^−1^ · v^−1^ versus -0.0270 m · s^−1^ · g^−1^ · v^−1^) suggests that the dissipation of braking is nearly counterbalanced by propulsion. It is conceivable that any difference must be further compensated for during the subsequent single support phase, most likely during the rebound ^8^.

While the collision impulse remains constant, the decrease in vertical push-off impulse with an increase in walking velocity suggests that humans experience a positive power deficit. This deficit appears to be compensated during the rebound phase ^8^. Additionally, there seems to be a tendency to minimize delayed push-offs, as indicated by the decrease in the total push-offs with higher walking velocities. Pre-emptive push-offs can help limit subsequent collisions ^7^. However, delayed push-offs or subsequent rebounds not only need to provide the necessary energy for the next stance but also must counteract the dissipation caused by a larger collision.

The increase in P/A braking and vertical GRF collision impulses with terrain amplitude may be linked to disturbances in relative collision and push-off timing ^18^. Consequently, sub-optimal push-offs may not fully restrict the collision magnitude, leading to the rise of collision impulses ^7,8^. The elevation of collision impulses with age and the interaction of age and terrain amplitude suggests that older adults, particularly on uneven terrains, may struggle more with efficiently modulating relative collision and push-off timing.

The increase in propulsive impulses beyond the braking phase suggests that humans utilize P/A impulses to propel forward during uneven walking ^25^. Normal lookahead aids in the targeting process for the next foot placement ^2^. Therefore, the regulation of P/A propulsive impulses may also contribute to setting the initial velocity of the COM.

The decrease in push-off impulses with increased terrain amplitude suggests a potential interruption in push-off generation. Push-off typically initiates just before the double support phase, predominantly by the ankle joint ^8^. As double support begins with the heel strike of the leading leg, heel strikes may interfere with the push-off effort of the trailing leg. Furthermore, the additional reduction in push-off with age and the interaction of age and terrain amplitude may indicate that older adults must compensate for larger collision dissipations. This scenario could also be exacerbated by disruptions in relative collision and push-off timing ^18^.

The interaction between age and terrain amplitude may be linked to their shared impact on gait. In both old-age and uneven walking conditions, deviations from typical walking patterns are observed. Muscle coactivations, where the agonist muscle generates a larger force to overcome the antagonist muscles for movement, are reported in both scenarios ^22,26^. Additionally, characteristics such as shorter step lengths, wider steps, and increased step variability are common in both old-age and uneven walking ^12,22,27,28^. While wider steps and increased variability assist in maintaining lateral balance and correcting steps in response to terrain perturbations [22], [29], [30], these same characteristics may also reflect deterioration in the Central Nervous System (CNS) function in older adults ^29^. The shorter step length observed in both cases, leading to a longer double support phase ^30^, could further indicate disruptions in relative collision and push-off timing.

We observed that variations in walking speed or terrain amplitude can affect GRF trajectories, especially the vertical component profile, causing deviations from nominal walking as changes in peak and trough amplitudes. As a result, it’s reasonable to speculate that when these skewed vertical GRF profiles are present we can expect elevated metabolic rates ^31^. Such skewed vertical GRF profiles are commonly observed in load-bearing situations and individuals with gait disorders ^15^.

The analysis of GRFs, while valuable, has its limitations and may not fully capture the dynamics of the walking system. For instance, in the case of vertical GRF collision impulses, despite their positive magnitude, they actually dissipate the mechanical energy of the COM ^8,20^. Therefore, it’s essential to adopt a more comprehensive approach that considers the work performed ^4,7,8^. Additionally, the definition of collision and push-off impulses of the vertical GRF is somewhat arbitrary (start and finish points), while the COM power trajectory during the stance phase is distinctly divided into four phases ^8^. Consequently, solely analyzing impulses exertions may not provide a clear understanding of their consequences.

In summary, our observations indicate that the P/A impulse is influenced by walking speed and terrain amplitude. In addition to the vertical GRF push-off impulse, the P/A propulsive impulse likely contributes to energizing the step-to-step transitions. As walking speed or terrain amplitude increases, the magnitude of the push-off impulse decreases, leading to an increase in the subsequent collision impulse. Hence, whereas the GRF analysis offers valuable insights into uneven walking, however, to understand more aspects of this complex locomotion pattern, additional analyses are essential.

## Acknowledgements

This work was supported in part by the Natural Sciences and Engineering Research Council of Canada (NSERC) Discovery and Canada Research Chair (Tier 1) programs.

## References

1. Muller B. Handbook of Human Motion. Springer Berlin Heidelberg; 2018.

2. Matthis JS, Fajen BR. Humans exploit the biomechanics of bipedal gait during visually guided walking over complex terrain. Proc R Soc B. 2013;280(1762):20130700. doi:10.1098/rspb.2013.0700

3. Masani K, Kouzaki M, Fukunaga T. Variability of ground reaction forces during treadmill walking. J Appl Physiol (1985). 2002;92(5):1885–1890. doi:10.1152/japplphysiol.00969.2000

4. Donelan JM, Kram R, Kuo AD. Mechanical work for step-to-step transitions is a major determinant of the metabolic cost of human walking. Journal of Experimental Biology. 2002;205(23):3717–3727. doi:10.1242/jeb.205.23.3717

5. Vaverka F, Elfmark M, Svoboda Z, Janura M. System of gait analysis based on ground reaction force assessment. Acta Gymnica. 2015;45(4):187–193. doi:10.5507/ag.2015.022

6. Keller TS, Weisberger AM, Ray JL, Hasan SS, Shiavi RG, Spengler DM. Relationship between vertical ground reaction force and speed during walking, slow jogging, and running. Clin Biomech (Bristol, Avon). 1996;11(5):253–259. doi:10.1016/0268-0033(95)00068-2

7. Kuo AD. Energetics of actively powered locomotion using the simplest walking model. J Biomech Eng. 2002;124(1):113–120. doi:10.1115/1.1427703

8. Kuo AD, Donelan JM, Ruina A. Energetic consequences of walking like an inverted pendulum: step-to-step transitions. Exerc Sport Sci Rev. 2005;33(2):88–97. doi:10.1097/00003677-200504000-00006

9. Schwartz MH, Rozumalski A, Trost JP. The effect of walking speed on the gait of typically developing children. J Biomech. 2008;41(8):1639–1650. doi:10.1016/j.jbiomech.2008.03.015

10. Laufer Y. Effect of age on characteristics of forward and backward gait at preferred and accelerated walking speed. J Gerontol A Biol Sci Med Sci. 2005;60(5):627–632. doi:10.1093/gerona/60.5.627

11. Kim WS, Kim EY. Comparing self-selected speed walking of the elderly with self-selected slow, moderate, and fast speed walking of young adults. Ann Rehabil Med. 2014;38(1):101–108. doi:10.5535/arm.2014.38.1.101

12. Yamada T, Maie K, Kondo S. The characteristics of walking in old men analysed from the ground reaction force. The Journal of Anthropological Society of Nippon. 1988;96(1):7–15. doi:10.1537/ase1911.96.7

13. Majlesi M, Farahpour N, Robertson GE. Comparisons of Spatiotemporal and Ground Reaction Force Components of Gait Between Individuals with Congenital Vision Loss and Sighted Individuals. Journal of Visual Impairment & Blindness. 2020;114(4):277–288. doi:10.1177/0145482X20940429

14. Oliveira CF, Vieira ER, Machado Sousa FM, Vilas-Boas JP. Kinematic Changes during Prolonged Fast-Walking in Old and Young Adults. Front Med. 2017;4:207. doi:10.3389/fmed.2017.00207

15. Aminiaghdam S, Rode C. Effects of altered sagittal trunk orientation on kinetic pattern in able-bodied walking on uneven ground. Biology Open. Published online January 1, 2017:bio.025239. doi:10.1242/bio.025239

16. Aminiaghdam S, Rode C, Müller R, Blickhan R. Increasing trunk flexion transforms human leg function into that of birds despite different leg morphology. J Exp Biol. 2017;220(Pt 3):478-486. doi:10.1242/jeb.148312

17. Müller R, Tschiesche K, Blickhan R. Kinetic and kinematic adjustments during perturbed walking across visible and camouflaged drops in ground level. Journal of Biomechanics. 2014;47(10):2286–2291. doi:10.1016/j.jbiomech.2014.04.041

18. Darici O, Temeltas H, Kuo AD. Optimal regulation of bipedal walking speed despite an unexpected bump in the road. Haddad JM, ed. PLoS ONE. 2018;13(9):e0204205. doi:10.1371/journal.pone.0204205

19. Voloshina AS, Kuo AD, Ferris DP, Remy CD. A Model-Based Analysis of the Mechanical Cost of Walking on Uneven Terrain. Bioengineering; 2020. doi:10.1101/2020.06.15.152330

20. Donelan JM, Kram R, Kuo AD. Simultaneous positive and negative external mechanical work in human walking. J Biomech. 2002;35(1):117–124. doi:10.1016/s0021-9290(01)00169-5

21. Shoja O, Farsi A, Towhidkhah F, Feldman AG, Abdoli B, Bahramian A. Visual deprivation is met with active changes in ground reaction forces to minimize worsening balance and stability during walking. Exp Brain Res. 2020;238(2):369–379. doi:10.1007/s00221-020-05722-0

22. Voloshina AS, Kuo AD, Daley MA, Ferris DP. Biomechanics and energetics of walking on uneven terrain. J Exp Biol. 2013;216(Pt 21):3963–3970. doi:10.1242/jeb.081711

23. Hunt AE, Smith RM, Torode M, Keenan AM. Inter-segment foot motion and ground reaction forces over the stance phase of walking. Clin Biomech (Bristol, Avon). 2001;16(7):592–600. doi:10.1016/s0268-0033(01)00040-7

24. Adamczyk PG, Kuo AD. Redirection of center-of-mass velocity during the step-to-step transition of human walking. J Exp Biol. 2009;212(Pt 16):2668–2678. doi:10.1242/jeb.027581

25. Franz JR. The Age-Associated Reduction in Propulsive Power Generation in Walking. Exercise and Sport Sciences Reviews. 2016;44(4):129–136. doi:10.1249/JES.0000000000000086

26. Lee HJ, Chang WH, Choi BO, Ryu GH, Kim YH. Age-related differences in muscle coactivation during locomotion and their relationship with gait speed: a pilot study. BMC Geriatr. 2017;17(1):44. doi:10.1186/s12877-017-0417-4

27. Owings TM, Grabiner MD. Step width variability, but not step length variability or step time variability, discriminates gait of healthy young and older adults during treadmill locomotion. J Biomech. 2004;37(6):935–938. doi:10.1016/j.jbiomech.2003.11.012

28. Brach JS, Berlin JE, VanSwearingen JM, Newman AB, Studenski SA. Too much or too little step width variability is associated with a fall history in older persons who walk at or near normal gait speed. J NeuroEngineering Rehabil. 2005;2(1):21. doi:10.1186/1743-0003-2-21

29. Bruijn SM, Van Dieën JH. Control of human gait stability through foot placement. J R Soc Interface. 2018;15(143):20170816. doi:10.1098/rsif.2017.0816

30. Kwon MS, Kwon YR, Park YS, Kim JW. Comparison of gait patterns in elderly fallers and non-fallers. Gómez C, Schwarzacher SP, Zhou H, eds. THC. 2018;26:427–436. doi:10.3233/THC-174736

31. O’Connor SM, Xu HZ, Kuo AD. Energetic cost of walking with increased step variability. Gait Posture. 2012;36(1):102–107. doi:10.1016/j.gaitpost.2012.01.014

